# DeepSynBa: Actionable Drug Combination Prediction with Complete Dose-Response Profiles

**DOI:** 10.1101/2025.01.28.634673

**Authors:** Halil Ibrahim Kuru, Haoting Zhang, Magnus Rattray, Carl Henrik Ek, A. Ercument Cicek, Oznur Tastan, Marta Milo

## Abstract

Many cancer monotherapies demonstrate limited clinical efficacy, making combination therapies a relevant treatment strategy. The extensive number of potential drug combinations and context-specific response profiles complicates the prediction of drug combination responses. Existing computational models are typically trained to predict a single aggregated synergy score, which summarises drug responses across different dosage combinations, such as Bliss or Loewe scores. This oversimplification of the drug-response surface leads to high prediction uncertainty and limited actionability, as these models fail to distinguish between potency and efficacy. We introduce DeepSynBa, an actionable model that predicts the complete dose-response matrix of drug pairs instead of relying on an aggregated synergy score. This is achieved by predicting parameters describing the response surface as an intermediate layer in the model. Evaluated on the NCI-ALMANAC and the O’Neil datasets, DeepSynBa outperforms the state-of-the-art methods in the dose-response matrix prediction task across most evaluation scenarios, including testing on novel drug combinations, cell lines, and drugs, across nine different tissue types. We also show that DeepSynBa yields reliable synergy score predictions. More importantly, DeepSynBa can predict drug combination responses across different dosages for untested combinations. The intermediate dose-response parameter layer enables the separation of efficacy from potency, informing the selection of dosage ranges that optimise efficacy while limiting off-target toxicity in experimental screens. The predictive capability and the downstream actionability make DeepSynBa a powerful tool for advancing drug combination research beyond the limitations of the current approaches. The code and the dataset for DeepSynBa are available at https://github.com/hikuru/DeepSynBa.

## Introduction

Over the past decade, high-throughput screening technologies have produced vast amounts of pharmacological data, including those from the DREAM challenge [Menden et al., 2019], NCI-ALMANAC [Holbeck et al., 2017], and the Wellcome Sanger Institute [Jaaks et al., 2022]. This progress has enabled data-driven approaches for preclinical modelling and the prediction of effective drug combination regimens.

Many frameworks have demonstrated progress in predicting the synergy scores of drug combinations using drug and cell line-related inputs such as transcriptomic profiles of the cell lines and the chemical structures of the drugs [Preuer et al., 2018, Kuru et al., 2021, El Khili et al., 2023, Wang et al., 2021a, Kuru et al., 2024, Jin et al., 2021, Bertin et al., 2023, Abbasi and Rousu, 2024]. Preuer et al. [2018] introduced DeepSynergy, in which the chemical structures and cancer cell line gene expression data are fed into a fully connected neural network to predict the Loewe synergy scores of drug combinations. Jin et al. [2021] designed a neural network architecture that jointly learns drug-target interactions and drug-drug synergies through a drug-target interaction module and a target-disease association module, which effectively identified synergistic drug combinations for treating COVID-19. Bertin et al. [2023] developed RECOVER, a sequential model optimisation search platform with a deep learning model to discover synergistic drug combinations. Kuru et al. [2021] designed MatchMaker that employs three fully connected neural networks to learn cell line-specific representations of drugs and predict the Loewe synergy scores of drug combinations. More recently, Kuru et al. [2024] developed PDSP, building upon MatchMaker by incorporating patient-specific single-drug response data to customise synergy predictions for individual patients.

Most drug synergy predictors, however, are trained with and predict a single synergy score, such as Loewe [Loewe, 1953] or Bliss [Bliss, 1939], which aggregates the drug combination effects across different dosage combinations into a single value. The reliance on a single summary score leads to several technical limitations that affect the models’ generalisability to real-world screening efforts. First, using a single synergy score can obscure different aspects of drug-drug interaction, such as potency and efficacy [Wooten et al., 2021]. Efficacy is the maximal effect a drug can achieve, whereas potency is the dose required for a drug to reach a specified effect [Rang et al., 2011]. In the PDSP model [Kuru et al., 2024], the potency of each drug (quantified by a binarised IC50) is separated, but efficacy is not. A second limitation is that there is no universally accepted gold standard for synergy scores [Tang et al., 2015]. Different synergy models, such as Loewe or Bliss, make different assumptions about drug interactions, leading to variations in the estimated synergy scores across studies [Vlot et al., 2019]. This variation reduces the generalisability of the models and compromises the use of these models in decision-making for drug discovery pipelines. Moreover, synergy scores are often associated with high uncertainty due to noise in drug-response measurements. Consequently, models based on synergy scores overfit a proxy measure that cannot estimate combination effects and is prone to high variability [Zhang et al., 2023]. Together, these limitations constrain the drug combination models’ ability to reliably predict drug combination effects.

In addition to technical limitations, the use of aggregated synergy scores for predicting combination effects also introduces experimental constraints that limit their applicability. In many screenings, dosage ranges are determined empirically with limited prior knowledge, often resulting in suboptimal coverage of dosages. If key dosages fall outside tested intervals, synergy scores can be misleading.

To address both modelling and experimental shortcomings, an emerging alternative is to predict the entire dose-response matrix rather than a single summary value. By predicting detailed response profiles across all tested concentration pairs, models can better characterise combination effects and calculate synergy scores as a secondary follow-up step. Few methods adopt this dose-response matrix prediction approach. For example, Julkunen et al. [2020] developed comboFM, which uses higher-order tensor factorisation to model drug interactions. Wang et al. [2021b] introduced comboLTR, which similarly uses polynomial regression on high-order tensors but replaces factorisation machines with latent tensor reconstruction. Huusari et al. developed comboKR [Huusari et al., 2025a] and its extension comboKR 2.0 [Huusari et al., 2025b], which use kernel-based functional regression to model drug combination dose-response surfaces. The latter extends the former with improved modelling choices and a projected gradient method, enhancing scalability and predictive performance. Rønneberg et al. [2023] proposed PIICM, which employs a permutation-invariant Gaussian process to predict the dose-response surfaces for untested drug combinations. Jin et al. [2025] introduced DD-PRiSM, a deep learning pipeline that decomposes the predicted dose-response matrix into monotherapy and synergistic components, where the monotherapy dose-response curves are pretrained from drug structure and cell line gene expression. A combination model is then built on top of the trained monotherapy models by the addition of weighted monotherapy contributions and a learned synergy term.

Although these methods advance beyond single-score prediction, they share a common limitation. ComboFM and comboLTR model the full response surface through latent factors without explicit pharmacological parameters. ComboKR, comboKR 2.0 and PIICM focus on predicting the overall response surface, without defining separate parameters for synergistic potency or synergistic efficacy. DD-PRiSM explicitly models monotherapy potency and efficacy, but for combinations, it only decomposes the predicted drug responses into monotherapy and synergy components, without introducing separate parameters for synergistic potency or synergistic efficacy. As a result, none of these methods explicitly evaluate synergistic potency and synergistic efficacy as separate quantities, which limits their utility in experimental screening workflows.

To address these technical and experimental limitations, we introduce DeepSynBa, a novel deep learning framework designed to predict the complete dose-response matrix of drug combinations on specific cell lines by disentangling the efficacy and potency of the combination. Fig. 1 illustrates the framework’s architecture. By training on the entire dose-response matrix, DeepSynBa captures comprehensive information across a range of dosages, offering a richer characterisation of the drug combination responses. Importantly, DeepSynBa explicitly incorporates distinct parameters for synergistic potency and synergistic efficacy, allowing separate learning of these critical aspects of a drug combination response.

**Figure 1.**
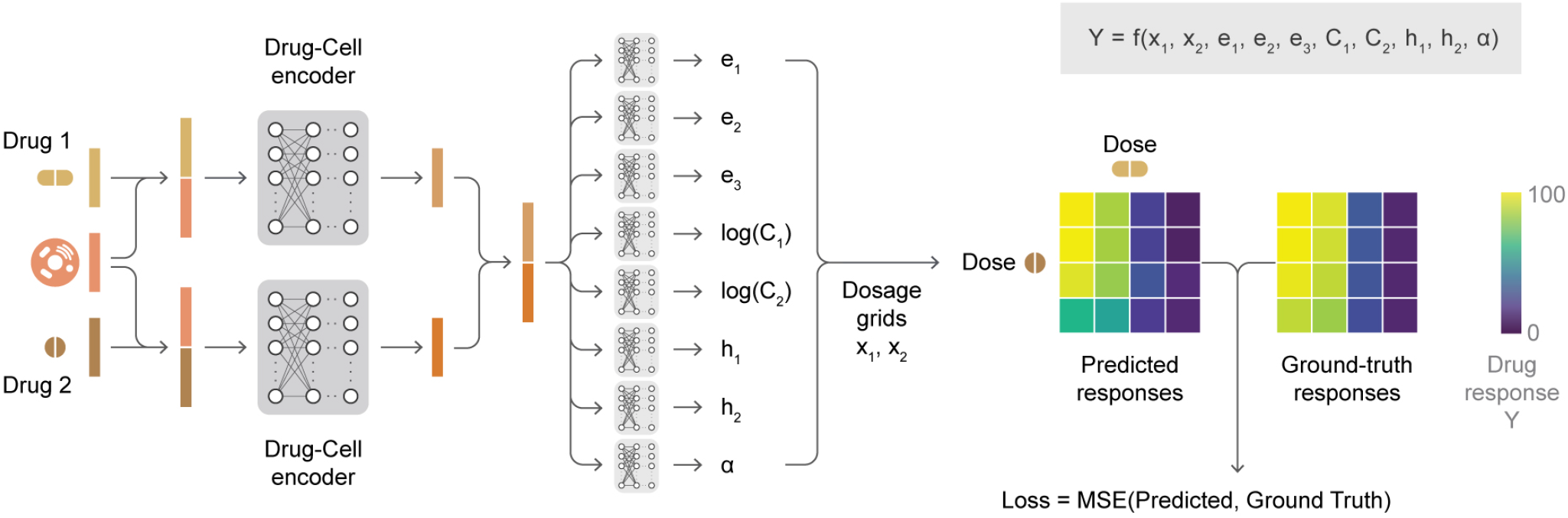
The general architecture of DeepSynBa. DeepSynBa aggregates drug and cell line features by concatenating them and passing the results through fully connected layers to obtain a unified representation. This unified representation is then fed into prediction heads to predict the eight parameters that characterise the dose-response surfaces. During training, the mean of the likelihood function from SynBa [Zhang et al., 2023] is used to predict the dose-response matrices from these parameters. The mean-squared error (MSE) loss is computed by comparing the predicted matrices with the ground truth ones.

We formulate the dose-response matrix prediction task as a regression problem, where the model is trained with dose-response matrices to predict matrix entries across a dosage grid. DeepSynBa employs the Matchmaker’s cell-line conditioned drug representation modules. To parameterise dose-response surfaces and disentangle the synergistic potency and efficacy, DeepSynBa follows the approach outlined in SynBa [Zhang et al., 2023] and adopts a simplified version of the two-dimensional generalised Hill Equation derived in MuSyC [Wooten et al., 2021]. DeepSynBa’s novel intermediate layer learns the parameters of the generalised Hill equation during training. Using these predicted parameters, the complete dose-response surface can be generated for a given drug combination as exemplified in Fig. 2 (a) and (e) panels. This model design also enables the computation of traditional synergy scores like Loewe and Bliss post-hoc, ensuring compatibility with existing drug synergy prediction methods while remaining flexible by not being constrained to any specific synergy score.

**Figure 2.**
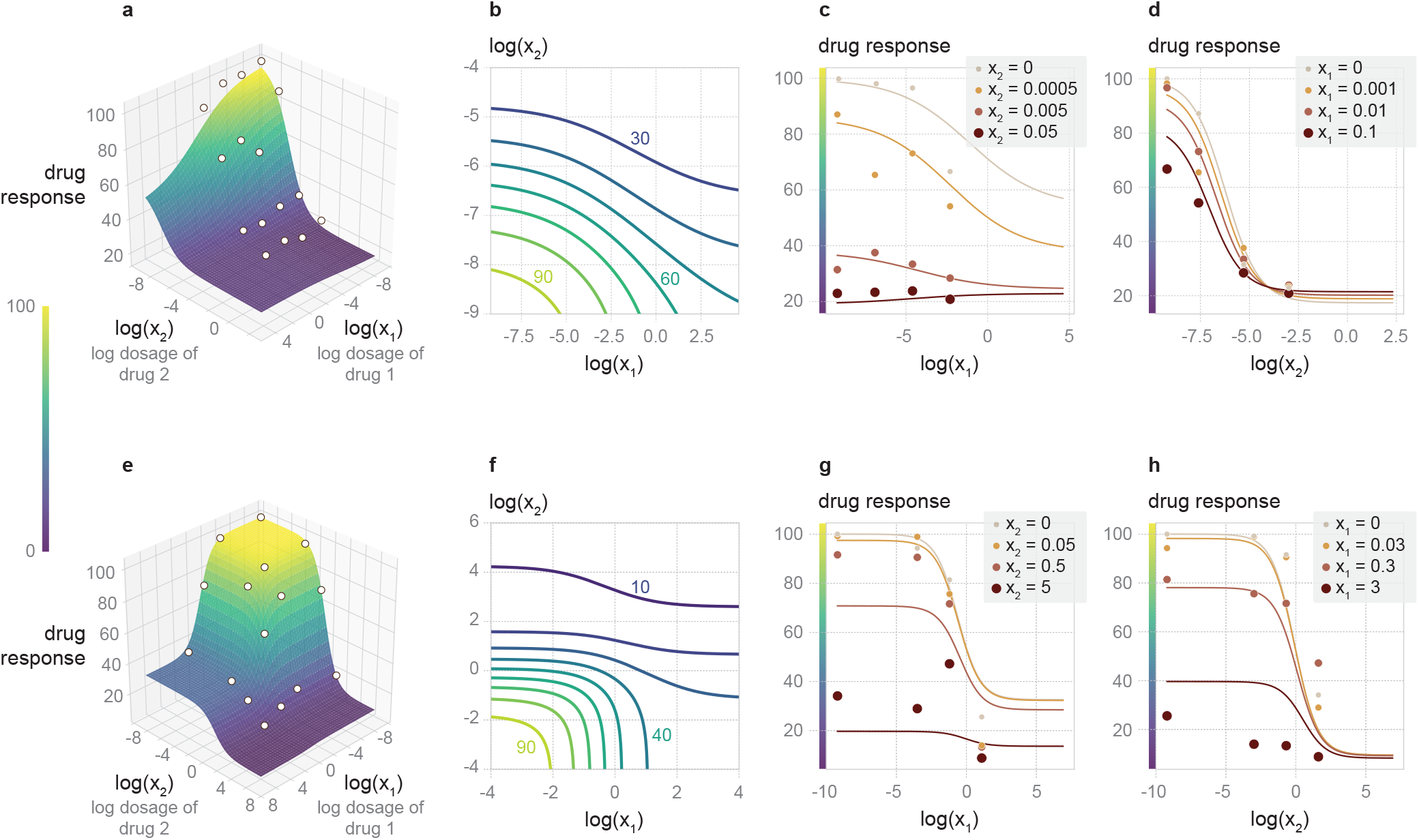
The predicted dose-response outputs from DeepSynBa for two different drug-cell line combinations and their comparisons with the ground-truth experimental measurements. (a)-(d) focus on the combination of Drug 1 (Bleomycin Sulfate) and Drug 2 (Cabazitaxel) administered on cell line 786-0. (e)-(h) correspond to the combination of Drug 1 (Azacitidine) and Drug 2 (Triethylenemelamine) administered on the cell line MDA-MB-468. (a)(e): Predicted dose-response surfaces and the measured responses (highlighted in white). (b)(f): Contour plots for the predicted drug response at various dosage levels of the two drugs. (c)(g): Dose-response curves for fixed concentrations of Drug 2 (in unit *μM*), comparing predictions with measurements (in *μM*). (d)(h): Dose-response curves for fixed concentrations of Drug 1 (in unit *μM*), comparing predictions with measurements (in *μM*).

Our contributions can be summarised as follows: DeepSynBa (i) provides a better dose-response prediction capability than existing dose-response predictors with its novel framework, (ii) allows separating out efficacy and potency which is not possible with existing models, and (iii) can assist experimental design by generating the dose-response surface across all dosages and thus identifying the suitable dosage range for the next batch of experiments.

## Methods

### Problem Formulation

We approach the dose-response prediction task as a regression problem, where we jointly predict the dose-response matrix elements. For each drug pair *< i, j >* and a cell line *k*, the target value is a matrix 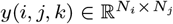, where *N*_*i*_ and *N*_*j*_ indicate the number of dosages of drugs *i* and *j*, respectively, used in the grid. Each drug *i* and *j* is represented by drug feature vectors *d*_*i*_ and *d*_*j*_, while the cell line *k* is represented by a cell feature vector *c*_*k*_. The goal is to predict the output drug responses 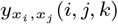 given the following inputs: the drug-specific features (*d*_*i*_ and *d*_*j*_), the cell line-specific features *c*_*k*_, and the dosages (*x*_*i*_, *x*_*j*_) used in the experiment.

### Dataset and Processing

We use the NCI-ALMANAC and the O’Neil dataset [Holbeck et al., 2017, O’Neil et al., 2016] to develop and evaluate our models and compare to other benchmarks. The NCI-ALMANAC dataset contains 274,516 drug combination matrices, covering 101 unique drugs and 60 unique cell lines. Each matrix in the dataset is 4 *×* 4, where the first row and first column correspond to single drug therapy responses. For the definition of the drug responses *y* in NCI-ALMANAC, we use the column `PercentGrowthNoTZ` (percent growth without time zero) instead of `PercentGrowth` (percent growth with time zero) in the original NCI-ALMANAC dataset. These two columns normalise the responses differently. In both columns, 100 represents the normalised number of cells in the control plate (no drug treatment). In `PercentGrowthNoTZ`, 0 means all cells are dead, ensuring that `PercentGrowthNoTZ` is non-negative, whereas in `PercentGrowth`, 0 means zero cell growth and −100 means all cells are dead. We opt for `PercentGrowthNoTZ` for two reasons. First, the generalised Hill equation is defined only for non-negative responses, which matches the range of `PercentGrowthNoTZ`. Second, in `PercentGrowth`, the positive numbers (representing cell growth) and negative numbers (between −100 and 0, representing cell death) do not share the same scale due to the way they are rescaled relative to the control plate. This makes it difficult to model `PercentGrowth` with a smooth function such as Equation 1.

The O’Neil dataset originally contains 21,361 dose-response matrices, covering 38 unique drugs and 39 unique cell lines across six tissue types. Each matrix is 4 *×* 4, but different from NCI-ALMANAC, all entries in O’Neil correspond to combination therapy responses at non-zero doses, with no single-agent monotherapy controls embedded in the matrix edges. In this study, ten cell lines (CAOV3, COLO320DM, DLD1, EFM192B, KPL1, LNCAP, MSTO, NCIH460, SKMEL30, ZR751) are excluded due to missing gene expression profiles in the DepMap database, and two drugs (L778123 and Oxaliplatin) are excluded due to unavailable molecular structure representations. After filtering, 14,031 matrices covering 36 drugs and 27 cell lines are retained.

For both datasets, for replicates measuring responses of the same combination and cell line at the same dose, we take the median value to handle the one-to-many mapping from the input space to the response.

### DeepSynBa Architecture

The proposed model architecture, illustrated in Fig. 1, is designed to predict the effects of drug combinations on cell lines by incorporating both drug and cell line features. Each drug combination consists of two drugs applied to a single cell line, and the model includes two dedicated drug-cell encoder sub-networks. Each encoder focuses on fusing the properties of one drug with the gene expression profiles of the cell line, enabling a detailed capture of interactions between the drugs and the cellular gene expression profiles.

The model uses feature representations for drugs (*d*_*i*_ and *d*_*j*_) and cell lines (*c*_*k*_). Drug features are generated as vector embeddings from the MoLFormer model [Ross et al., 2022], and the cell line features are represented by vectors of untreated gene expression data. These features are concatenated into a combined vector ([*d*_*i*_; *c*_*k*_] or [*d*_*j*_ ; *c*_*k*_]) and then processed through a series of fully connected layers. These layers transform the combined drug-cell line features into a comprehensive representation that captures the interactions between the drugs and the cell line. The outputs of the drug-cell encoders are gene expression-based feature maps, which encapsulate the interaction between each drug and the cell line at a molecular level. The feature maps from each of the encoders are then concatenated to produce a unified representation, effectively summarising the combined impact of both drugs on the cell line.

Subsequently, the unified representation is input into eight distinct prediction heads, each corresponding to one parameter in the SynBa likelihood function [Zhang et al., 2023]. Each prediction head comprises fully connected layers that process the unified representation to predict a single scalar value for each parameter. By using these prediction heads, the model can estimate the SynBa parameters accurately.

Specifically, the prediction heads are *e*_1_, *e*_2_, *e*_3_, *C*_1_, *C*_2_, *h*_1_, *h*_2_ and *α* from the likelihood function defined in SynBa, which are the parameters that characterise the expected value of the drug responses *y*. Given the dosages *x*_1_ for Drug 1 and *x*_2_ for Drug 2, the expected value for a response *y* is defined as

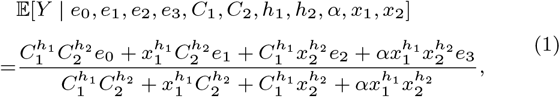

where *e*_0_ is the base value for the drug effect when no drug is applied, *e*_1_, *C*_1_, and *h*_1_ are the monotherapy parameters associated with Drug 1, *e*_2_, *C*_2_, and *h*_2_ are the monotherapy parameters associated with Drug 2, and *e*_3_ is the maximal effect for the combined drugs. The association parameter *α* controls how the two drugs are affected by the presence of each other, where *α >* 1 is equivalent to *synergistic potency*, meaning a smaller amount of drug is required to reach the same level of dosage when the two drugs are combined. For the stability of training, *e*_0_ is set to be a fixed value in DeepSynBa and determined by the definition of the dataset. For example, the base value is 100 in NCI-ALMANAC.

Our approach leverages the mean function of the Gaussian likelihood for drug responses from SynBa [Zhang et al., 2023], given in Equation 1. This function stems from the two-dimensional generalised Hill Equation derived in MuSyC [Meyer et al., 2019], in which the classical Hill equation

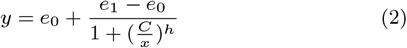

for monotherapy was extended to incorporate the combination of two drugs. Among the five levels of complexity in the nested structure of the two-dimensional generalised Hill Equation in MuSyC, we adopt the most complex tier that still assumes detailed balance, the same as the choice in SynBa [Zhang et al., 2023]. This selection balances accurately modelling the biologically viable dose-response surfaces and avoiding overparameterisation. The latter problem, if not handled, would raise the problem of non-identifiability in the deep networks. The outputs from the eight prediction heads are then input into the SynBa likelihood function to compute the dose-response matrices. By embedding the SynBa likelihood function within the prediction pipeline, the model can be trained end-to-end and can predict dose-response surfaces that reflect the effects of varying drug doses on the cell line response.

### Training details

We evaluate various hyperparameter settings to train DeepSynBa. The Drug-Cell encoder consists of a single hidden layer, with layer sizes from *{*256, 512, 1024, 2048, 4096*}*, followed by batch normalisation, ReLU activation, and dropout. The model includes eight prediction heads, each containing three blocks. The input layer size of the prediction head is chosen from *{*64, 128, 256*}*, and each block progressively reduces the size to one-fourth of the previous layer. Each block contains a fully connected layer, followed by layer normalisation, ReLU activation, and dropout. The final layer applies either a sigmoid or linear activation based on the range of the corresponding SynBa likelihood function parameter. Training is conducted for up to 500 epochs using the AdamW optimiser, with a learning rate of 10^−4^, weight decay of 10^−5^, and a batch size of 128. All hyperparameters are tuned on the validation data minimising loss. The details are available in Supplementary Table S3.

We train all the models on a SuperMicro SuperServer 4029GB-TRT with 2 Intel Xeon Gold 6140 Processors (2.3GHz, 24.75M cache) and 251GB RAM. The models are trained on 1 NVIDIA TITAN RTX GPU (24GB, 384Bit).

## Results

### Case Studies

To demonstrate how DeepSynBa provides deeper insights by predicting the full dose-response matrix or surface, we examine two specific examples in the NCI-ALMANAC dataset. The first example focuses on the combination of Bleomycin Sulfate (denoted as Drug 1) and Cabazitaxel (denoted as Drug 2) administered on the renal cancer cell line 786-0. Drug 1 is an antibiotic with a chemotherapy effect on fast-growing cells, while Drug 2 is a strong chemotherapy agent used in the treatment of prostate cancer.

Fig. 2(a)-(d) illustrates the predicted and measured effects of the drug combination. Panel (a) presents the predicted dose-response surface, with the actual measured responses coloured in white. The surface predicted by DeepSynBa matches the downward pattern of the measured responses as the dosages of both drugs increase. The killing effect of the drugs is quantified as a drop along the y-axis.

Panel (b) presents the corresponding contour plot for the dose-response surface, highlighting the combinations of dosage levels that reach the predicted drug effects. The plot reveals a rapid decline along the y-axis from 100% to 60%, indicating fast cell killing as the dosages for both drugs increase. More interestingly, this drop is accompanied by evidence of synergistic potency, as indicated by the curvature of the contour lines for response levels 90%, 80%, 70%, and 60%. For a combination that is neither synergistic nor antagonistic, the contour lines for dose-response levels typically resemble a circle or an oval. In panel (b), the curvature bends inward towards the origin, showing that a smaller dosage is required when the drugs are combined to achieve the same level of efficacy as single-drug treatments. However, beyond the 60% response level, increasing the dosage of Drug 1 (right side of the x-axis) does not significantly improve the treatment response. In contrast, Drug 2 continues to exhibit increasing effectiveness as the dosage increases. This aligns with the fact that Drug 2 is a strong chemotherapy agent, and it is expected to become dominant at higher dosage levels for both drugs.

Panel (b) also demonstrates that even for the same drug combination, the synergy profile can vary across different dosage ranges. Synergistic potency may only be present within specific dosage subsets. This further highlights the importance of predicting the full dose-response surface to identify actionable combinations and their respective dose ranges.

The ability to estimate the full drug combination profile surface allows us to take slices of the surface for further inspection of the drug’s interaction and effects. For panels (c) and (d), slices were taken from the dose-response surface at the dosage levels where actual measurements were available, enabling an evaluation of predictions against the observed data. The measured responses (represented with the dots) and predicted dose-response curves are shown for each of the two drugs.

In both panels (c) and (d), it can be observed that at lower doses of one drug, the curves shift downward as the dose of the other drug increases. However, when both drugs are administered at higher doses, the difference between the curves narrows. This is particularly evident in (d), where the predicted maximal efficacy does not improve as the dose of Drug 1 increases. Nevertheless, its drug response reaches the IC50 at a lower dosage when combined with the other drug. This shows the potential benefit of combining these two drugs at a medium-to-low dose range. Such a combination could reduce the risk of off-target toxicity associated with high doses of the strong chemotherapy agent, achieving the desired therapeutic effect with lower doses of each drug compared to their monotherapy counterparts.

In the second example, we look at the combination of Azacitidine (Drug 1) and Triethylenemelamine (Drug 2), two strong chemotherapy agents, administered on MDA-MB-468, a triple-negative breast cancer cell line. Azacitidine is used in leukaemia, while Triethylenemelamine is particularly effective against rapidly growing cancer cells. In Fig. 2, panels (e)-(h) illustrate their predicted behaviour and the comparisons with the actual measurements. In panel (f), it is evident that there is no clear synergistic interaction between these two drugs, as the contour lines from 80% to 40% mostly bend outwards, away from the origin (i.e. the bottom-left corner of the plot). Despite both drugs being effective on MDA-MB-468, combining them does not offer a clear advantage for this particular tumour indication. This is an example showing how our model helps to identify the lack of beneficial interactions in a specific setting, providing insight into why certain combinations might not be suitable for further testing in that tumour type.

These two examples demonstrate the ability of DeepSynBa to model and predict how two drugs interact while separately quantifying two key combination metrics: synergistic potency and maximal efficacy. This approach provides not only clearer biological insights into their combined effect but also valuable guidance for decision-making when evaluating drug combination dose regimes.

Fig. 2 illustrates clearly that the analysis performed in the previous sections would not be possible if the entire combination profile were summarised into a single synergy score. This richer information provided by the full-dose response matrix is crucial for optimising combination therapies, as it identifies regions in the dose space where drugs can achieve significant benefit through enhanced potency without increasing their maximum effect. This could help guide experimental design in collecting further evidence for the combination regimes and ultimately supporting the development of safer and more effective treatment strategies.

### Performance Evaluation Setup

We evaluate the performance of our models across three stratification scenarios: (i) new drug combinations, (ii) new cell lines, and (iii) new drugs. In the case of new drug combinations, a specific drug pair *< i, j >* is restricted to appearing in only one of the three datasets: training, validation, or testing. For the new cell scenario, drug combinations are grouped by cell line types, ensuring that any given cell line *k* is included exclusively in one of the splits - training, validation, or test. In the new drug scenario, drug combinations in the test set must include at least one drug that is absent from any combinations used in the training or validation sets. Similarly, the validation set must contain drug combinations with at least one drug not present in the training set. In all three scenarios, the data is split into train, validation and test with 60%, 20% and 20%, respectively. These distinct evaluation strategies are critical for accurately assessing the predictive performance of the models in novel contexts. By structuring our evaluation in this way, we simulate real-world conditions where the introduction of new drugs and cell lines is possible, thereby providing a robust framework to assess the utility of our models in drug discovery and personalised medicine.

### Compared Methods

We evaluated our model’s performance against comboFM [Julkunen et al., 2020], comboLTR [Wang et al., 2021b], comboKR 2.0 [Huusari et al., 2025b] and DD-PRiSM [Jin et al., 2025], four models designed for predicting dose-response outcomes in drug combination studies. They are the most similar to DeepSynBa in terms of the data modalities used and the outputs predicted. For a fair comparison, we utilised the inherent drug and cell line feature representations of each baseline as originally proposed, without any substitution. DD-PRiSM is additionally provided with the monotherapy measurements in the O’Neil data source, which are distinct from the dose-response matrices, to fulfil its monotherapy pretraining stage.

While PIICM also predicts dose-response surfaces, its primary focus is on modelling correlations between dose-response functions across different experiments. Cell or drug properties are not incorporated, restricting its scope to cell lines and drugs that have already been tested. Therefore, it cannot be directly applied to evaluation scenarios which involve new drugs and new cell lines, so we did not include it in our experiments.

### Drug-Response Matrix Prediction Performance

To comprehensively evaluate models’ performances, we assess each model across various scenarios using three core metrics: root mean squared error (RMSE), Pearson correlation, and Spearman correlation. These metrics quantify the alignment between predicted dose-response matrices and the ground-truth values. The metrics are calculated over all entries of the dose-response matrices. Table 1 summarises each model’s performance in terms of RMSE, Pearson, and Spearman correlations across three scenarios and two datasets. We have excluded comboLTR from the O’Neil comparison because our training for comboLTR could not produce stable results on O’Neil, despite following the original implementation.

**Table 1.** Performance comparison of all methods in three different scenarios: (i) new drug combination, (ii) new cell line, and (iii) new drug, evaluated on two datasets: NCI-ALMANAC and O’Neil. These scenarios are described in Section 3.2.

|  | (i) New drug combination |  |  | (ii) New cell line |  |  | (iii) New drug |  |  |
| --- | --- | --- | --- | --- | --- | --- | --- | --- | --- |
|  | RMSE | Pearson | Spearman | RMSE | Pearson | Spearman | RMSE | Pearson | Spearman |
| NCI-ALMANAC |  |  |  |  |  |  |  |  |  |
| comboFM | 12.76 | 0.87 | 0.68 | 20.03 | 0.72 | 0.56 | 42.39 | 0.35 | 0.24 |
| comboLTR | 16.11 | 0.78 | 0.64 | 49.30 | 0.26 | 0.23 | 40.22 | 0.40 | 0.40 |
| comboKR 2.0 | 23.86 | 0.65 | 0.55 | 23.29 | 0.63 | 0.51 | 25.79 | 0.60 | 0.53 |
| DD-PRISM | 13.61 | 0.88 | 0.73 | 23.74 | 0.51 | 0.49 | 21.82 | 0.60 | 0.48 |
| DeepSynBa | **7.72** | **0.95** | **0.82** | **12.57** | **0.85** | **0.69** | **18.69** | **0.71** | **0.64** |
| O’NEIL |  |  |  |  |  |  |  |  |  |
| comboFM | 0.33 | 0.65 | 0.68 | 0.32 | 0.67 | 0.68 | 0.94 | 0.14 | 0.36 |
| comboKR 2.0 | 0.11 | 0.95 | 0.95 | **0.11** | **0.95** | **0.95** | **0.11** | **0.94** | **0.94** |
| DD-PRISM | 0.17 | 0.86 | 0.87 | 0.32 | 0.49 | 0.48 | 0.33 | 0.36 | 0.34 |
| DeepSynBa | **0.09** | **0.96** | **0.96** | 0.15 | 0.90 | 0.90 | 0.31 | 0.53 | 0.55 |

In particular, DeepSynBa consistently outperforms all competing models in each experimental setting, except for in Scenarios (ii) and (iii) of the O’Neil dataset when compared to comboKR 2.0.

On NCI-ALMANAC, DeepSynBa achieves RMSE values of 7.72, 12.57, and 18.69 in Scenarios (i), (ii), and (iii) respectively, representing substantial margins over the next-best model in each case. For the O’Neil dataset, DeepSynBa achieves RMSE values of 0.09, 0.15, and 0.31 for the three scenarios. Note that the difference in RMSE scales between the two datasets reflects their different response normalisation conventions. Responses for control plates in NCI-ALMANAC are scaled to 100, whereas in O’Neil they are scaled to 1. While all models apart from comboKR 2.0 experience performance degradation in the more challenging Scenarios (ii) and (iii), DeepSynBa maintains its advantage across all three metrics except for comboKR 2.0 in Scenarios (ii) and (iii). This result suggests the effectiveness of DeepSynBa’s architecture in accurately predicting drug combination responses.

We note that comboKR 2.0 is the only method that shows no significant performance drop from Scenario (i) to (ii) and (iii) across both datasets. Unlike other methods, it constructs full kernel matrices over all entities prior to training, meaning the features of test cell lines and drugs are encoded in the kernel matrices during training. In high-throughput screenings like NCI-ALMANAC and O’Neil where drug or cell line responses are often systematically correlated and biased due to shared experimental conditions [Caraus et al., 2015], this may give the method an advantage in the harder scenarios.

### Performance on Cell Lines and Drugs

We also calculated the performance metrics separately for each cell line and drug, enabling a detailed view of model performance across various tumour indications and drug pharmacokinetics. We focus on Scenario (i) with NCI-ALMANAC for this and the following stratified analyses, as NCI-ALMANAC covers substantially more drugs and cell lines than O’Neil, and Scenario (i) retains the largest proportion of these for evaluation compared to Scenarios (ii) and (iii).

These results are presented as distributions of performance metrics obtained for cell lines and drugs in Fig. 3. A pairwise t-test is applied to evaluate the statistical significance of performance differences between methods. All pairwise comparisons are statistically significant, with *p*-values *<* 10^−10^ (detailed results are provided in Supplementary Tables S1 and S2). As shown in Fig. 3, the RMSE distributions for cell line-based and drug-based results are concentrated at lower values for DeepSynBa, while the Pearson and Spearman correlation distributions are concentrated at higher values. These concentrated distributions reflect DeepSynBa’s superior predictive performance. Results from the t-test confirm that these performance improvements are statistically significant, highlighting DeepSynBa’s advantage in predicting dose-response outcomes. Moreover, the shorter height of DeepSynBa’s violin plots suggests its performance is consistent across different cell lines and drugs, indicating greater robustness.

**Figure 3.**
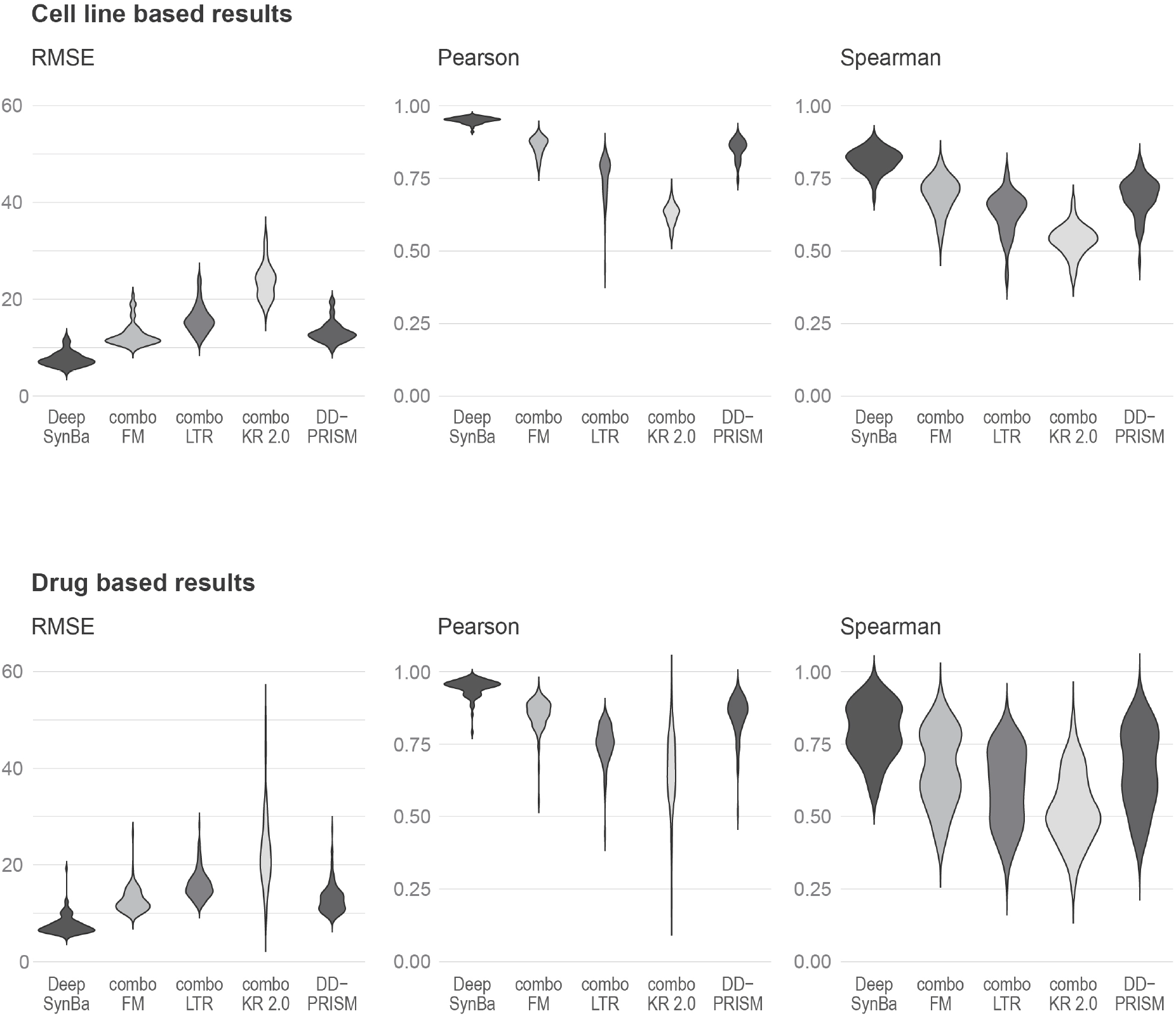
The distributions of RMSE, Pearson and Spearman correlations of dose-response surface predictions on different cell lines (upper panel) and drugs (lower panel) in Scenario (i) (new drug combinations) for training on the NCI-ALMANAC dataset.

### Performance Across Tissues

We also evaluated model performance across tissue types. The cell lines are grouped by their tissue of origin, and performance metrics are calculated within each group to assess variations in predictive accuracy across different tissues. Although performance improvements vary across tissues (see Table 2), DeepSynBa consistently outperforms other models across all examined tissue types. The model achieves the highest predictive accuracy for dose-response predictions in renal, breast, and ovarian cancers while exhibiting lower performance in leukaemia. This discrepancy likely arises from the distinct biological characteristics of these cancers and the nature of the screening assays, which tend to be more effective on solid tumours. Breast and ovarian cancers are characterised by well-defined genetic markers, such as BRCA1/2 mutations, which may facilitate model learning by providing clearer molecular patterns [Radu et al., 2021, Barili et al., 2024]. Similarly, renal cancers are often associated with specific signalling pathways, such as PI3K/AKT/mTOR, which are targets for therapeutic interventions and support more reliable prediction of therapeutic responses [Miricescu et al., 2021, Guo et al., 2015]. In contrast, leukaemia lacks a solid tumour structure and displays significant genetic heterogeneity, posing challenges for predictive models [Tang et al., 2023, Estey et al., 2015, Calderon et al., 2024].

**Table 2.** Performances based on tissue types in Scenario (i) (new drug combinations) for training on the NCI-ALMANAC dataset.

|  | RMSE |  |  |  |  | Pearson |  |  |  |  |
| --- | --- | --- | --- | --- | --- | --- | --- | --- | --- | --- |
|  | DeepSynBa | comboFM | comboLTR | comboKR | DD-PRISM | DeepSynBa | comboFM | comboLTR | comboKR | DD-PRISM |
| Breast | **6.67** | 11.39 | 14.05 | 21.57 | 12.36 | **0.95** | 0.86 | 0.77 | 0.63 | 0.87 |
| CNS | **7.14** | 12.35 | 14.20 | 23.73 | 12.48 | **0.96** | 0.87 | 0.81 | 0.63 | 0.89 |
| Colon | **8.55** | 13.48 | 17.25 | 25.45 | 14.89 | **0.95** | 0.87 | 0.77 | 0.63 | 0.87 |
| Leukemia | **11.09** | 18.62 | 22.34 | 31.08 | 19.33 | **0.96** | 0.87 | 0.80 | 0.68 | 0.88 |
| Melanoma | **7.30** | 11.76 | 15.49 | 23.80 | 13.12 | **0.95** | 0.87 | 0.75 | 0.62 | 0.87 |
| Non-Small Cell Lung | **7.15** | 11.76 | 15.11 | 22.39 | 12.63 | **0.95** | 0.87 | 0.77 | 0.64 | 0.88 |
| Ovarian | **6.97** | 11.76 | 15.46 | 21.40 | 12.30 | **0.95** | 0.85 | 0.72 | 0.64 | 0.87 |
| Prostate | **7.09** | 11.73 | 15.09 | 23.12 | 12.03 | **0.96** | 0.88 | 0.78 | 0.64 | 0.90 |
| Renal | **6.93** | 11.16 | 14.87 | 21.85 | 12.03 | **0.95** | 0.87 | 0.75 | 0.63 | 0.88 |

### Performance Across Drug Categories

We further analysed the results by categorising the drugs in the combinations according to NCI-ALMANAC’s classification: chemotherapy, targeted, and other. For two-drug combinations, six distinct pairings emerge from these categories. Performance metrics are computed separately for these pairings. Across all combinations, our model, DeepSynBa, consistently outperforms the competing methods, as demonstrated in Table 3. The most accurate dose-response predictions are observed when one drug belongs to the “other” category, paired with either a “targeted” drug or another “other” drug. This phenomenon can likely be attributed to the distribution of responses within these categories. Specifically, the “Targeted-Other” and “Other-Other” combinations show minimal impact on the overall response, as their distributions are concentrated within a narrow range (see Supplementary Fig. S3), making the prediction task more tractable.

**Table 3.** Performances based on drug combination categories in Scenario (i) (new drug combinations) for training on the NCI-ALMANAC dataset.

|  | RMSE |  |  |  |  | Pearson |  |  |  |  |
| --- | --- | --- | --- | --- | --- | --- | --- | --- | --- | --- |
|  | DeepSynBa | comboFM | comboLTR | comboKR | DD-PRISM | DeepSynBa | comboFM | comboLTR | comboKR | DD-PRISM |
| Chemotherapy-Chemotherapy | 8.07 | 13.03 | 16.57 | 25.84 | 14.30 | **0.96** | 0.88 | 0.80 | 0.64 | 0.89 |
| Chemotherapy-Targeted | 8.33 | 12.39 | 16.16 | 20.61 | 14.95 | **0.94** | 0.86 | 0.73 | 0.66 | 0.87 |
| Chemotherapy-Other | 7.29 | 13.48 | 16.23 | 25.37 | 12.91 | **0.96** | 0.86 | 0.79 | 0.65 | 0.89 |
| Targeted-Targeted | 6.32 | 11.77 | 15.55 | 13.10 | 13.95 | **0.94** | 0.79 | 0.54 | 0.71 | 0.81 |
| Targeted-Other | 5.69 | 10.45 | 12.95 | 16.62 | 11.16 | **0.94** | 0.81 | 0.68 | 0.66 | 0.87 |
| Other-Other | 5.77 | 11.29 | 14.69 | 26.80 | 9.19 | **0.97** | 0.88 | 0.79 | 0.65 | 0.90 |

### Synergy Score Prediction Performance

To assess compatibility with existing synergy score frameworks, we derive Loewe and Bliss scores from the predicted dose-response matrices of NCI-ALMANAC and compare them against ground-truth values. The results, presented in Table 4, demonstrate that DeepSynBa consistently outperforms the other drug combination matrix prediction methods in terms of each metric. These findings highlight the superior performance of DeepSynBa relative to alternative approaches in capturing drug synergy, providing robust evidence for its efficacy in predicting synergistic interactions.

**Table 4.**
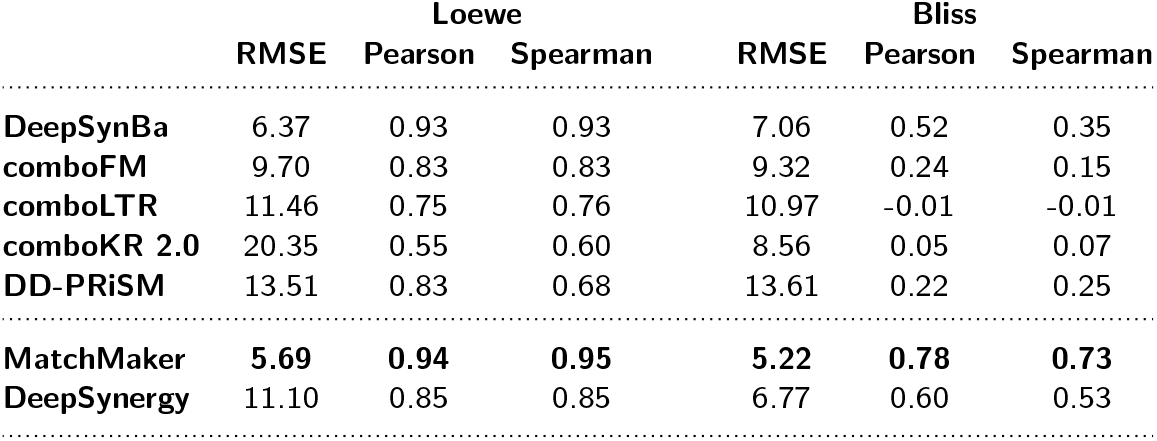
Performances with the prediction of Loewe and Bliss synergy scores in Scenario (i) (new drug combinations) for training on the NCI-ALMANAC dataset. Ground-truth Loewe/Bliss scores are calculated with dose-response matrices. Predicted Loewe/Bliss scores for DeepSynBa, comboFM and comboLTR models are calculated with predicted dose-response matrices. MatchMaker and DeepSynergy models are directly trained with Loewe/Bliss synergy scores.

|  | Loewe |  |  | Bliss |  |  |
| --- | --- | --- | --- | --- | --- | --- |
|  | RMSE | Pearson | Spearman | RMSE | Pearson | Spearman |
| **DeepSynBa** | 6.37 | 0.93 | 0.93 | 7.06 | 0.52 | 0.35 |
| **comboFM** | 9.70 | 0.83 | 0.83 | 9.32 | 0.24 | 0.15 |
| **comboLTR** | 11.46 | 0.75 | 0.76 | 10.97 | -0.01 | -0.01 |
| **comboKR 2.0** | 20.35 | 0.55 | 0.60 | 8.56 | 0.05 | 0.07 |
| **DD-PRISM** | 13.51 | 0.83 | 0.68 | 13.61 | 0.22 | 0.25 |
| **MatchMaker** | **5.69** | **0.94** | **0.95** | **5.22** | **0.78** | **0.73** |
| **DeepSynergy** | 11.10 | 0.85 | 0.85 | 6.77 | 0.60 | 0.53 |

We also assess the performance of two synergy score predictor models, MatchMaker [Kuru et al., 2021] and DeepSynergy [Preuer et al., 2018], which directly learn synergy scores. Although DeepSynBa is not optimised to predict Loewe or Bliss synergy scores directly, it achieves comparable synergy score prediction performance, especially in the case of Loewe scores. This result shows DeepSynBa’s capacity to generalise to downstream tasks without further task-specific supervised training, reducing the risk of overfitting.

### Decoupled prediction through intermediate parameter layer

In this subsection, we examine how the DeepSynBa intermediate layer enables decoupled prediction of synergistic potency and efficacy for previously unseen drug combinations. We focus on Scenario (i) of the NCI-ALMANAC setting, corresponding to the results reported in Table 1(i). Since synergistic potency and efficacy are not directly observed but can be inferred from dose-response matrices, we use SynBa [Zhang et al., 2023] as a proxy for ground truth. SynBa, as a Bayesian inference method, infers the full posterior distributions of all parameters in the 2-dimensional generalised Hill equation given a full dose-response matrix.

We fit SynBa to the full dose-response matrices and extract posterior mean estimates of synergistic potency, maximal efficacy, and synergistic efficacy (min{*e*_1_, *e*_2_} − *e*_3_). The synergistic potency (*α*) and maximal efficacy (*e*_3_) are parameters of the two-dimensional generalised Hill equation, as described in Section 2.3. The synergistic efficacy, defined as min*{e*_1_, *e*_2_*}*−*e*_3_ and denoted ΔHSA, quantifies the gain in maximal effect relative to the more efficacious monotherapy, with positive values indicating a benefit beyond what either drug achieves alone. We then compare these inferred parameters to the corresponding predictions produced by the DeepSynBa intermediate layer on the held-out test combinations.

Figures S4 and S5 show the correlation between the posterior mean estimates and the corresponding out-of-sample predictions for synergistic potency, maximal efficacy, and synergistic efficacy, respectively. The first row of the figures report the overall correlation across all 55,120 held-out drug combination instances, where each point corresponds to a single drug combination tested on a specific cell line. Across the held-out combinations, we observe Pearson correlations of 0.28, 0.43, and 0.36 for these three quantities that describe different aspects of a combination. This level of correlation indicates that the intermediate layer captures transferable signals for each component.

In contrast to scalar synergy prediction, the present setting requires simultaneous prediction of multiple mechanistically distinct surface parameters. Synergistic potency, maximal efficacy, and synergistic efficacy emerge jointly from the learned representation. The correlation across these distinct endpoints provides evidence that the intermediate layer has learned a meaningful and disentangled representation of drug combination effects, supporting its role as an interpretable and mechanistically grounded intermediate layer.

Beyond the overall correlations, Figures S4 and S5 reveal a visible discrepancy between the distributions of inferred and predicted parameter values. The inferred values are substantially more concentrated than the predictions for synergistic potency and maximal efficacy. To investigate this further, Fig. S6 bins the inferred parameters and shows the distribution of predictions within each bin. The median predicted value follows a broadly increasing trend across bins for all three parameters, suggesting the model captures relative ordering. However, the predicted distributions within each bin are wide, indicating that the model cannot consistently discriminate between individual instances within a given range. This reflects a challenge in disentangling surface parameters from the dose-response matrices, and motivates the probabilistic extension discussed in future work.

### Effect of Input Feature Modalities on Performance

We evaluated alternative representations for both drugs and cell lines to see the effect of different feature representations. For drug features, we compared MoLFormer embeddings with ECFP4 fingerprints in the new drug combination setting. The results are presented in Supplementary Table S4. While ECFP4 shows a slight performance advantage, MoLFormer achieves highly comparable results, indicating that transformer-based representations effectively capture relevant chemical information from SMILES strings.

For cell line features, we extended the original gene expression (GEX) representation by incorporating mutation (MUT), copy number variation (CNV), and DNA methylation (METHY) data into a unified feature vector. As shown in the Supplementary Table S5, the extended representation does not improve performance over GEX alone, suggesting that gene expression remains the most informative modality in this setting.

### Effect of Prediction Head Architecture

To evaluate the impact of the prediction strategy, we replaced the original parameter prediction head with a direct matrix prediction approach. Instead of estimating SynBa parameters and reconstructing the dose-response surface, the model predicts a 4 *×* 4 response matrix *R*_*i*_ for each drug. The final response is computed as a weighted combination of these matrices using the corresponding dosages (*d*_1_, *d*_2_). As shown in the Supplementary Table S6, DeepSynBa with parameter prediction substantially outperforms the matrix prediction head across all metrics, indicating the effectiveness of modelling underlying dose-response parameters rather than directly predicting responses.

## Discussion and Conclusions

In this study, we propose DeepSynBa which predicts full dose-response matrices of drug combinations while disentangling efficacy and potency. The model also enables the prediction of full dose-response surfaces. We assessed the performance of our model under three evaluation scenarios: (i) previously unseen drug combinations, (ii) new cell lines, and (iii) new drugs. In all settings with the exception of comboKR 2.0 in Scenarios (ii) and (iii) for the O’Neil dataset, DeepSynBa outperforms existing models in predicting dose-response matrices.

One limitation of our model is in the prediction of previously unseen new cells and drugs. Table 1(iii) shows RMSE and correlation metrics substantially degrade compared to the first scenario, Table 1(i). In the new cell scenario, performance falls in the mid-range. While it declines compared to the new drug combination setting, it still surpasses the performance observed in the new drug scenario. This suggests that while DeepSynBa partially utilises cell line specificity, further improvements are needed to enhance its ability to learn drug characteristics to generalise better across previously unseen drugs.

The reduced performance observed when predicting responses for previously unseen drugs reflects a broader open challenge in the field. Generalising to novel compounds is inherently difficult because the model must extrapolate from chemical and biological patterns that may not have been observed during training. Recent work [Çandır et al., 2026] has shown that simple representations such as one-hot drug encodings can perform competitively when compared with more sophisticated molecular embeddings in certain drug response prediction settings, suggesting that current molecular representations may not always capture the full range of pharmacologically relevant information required for reliable extrapolation to unseen compounds.

By developing DeepSynBa, we seek to encourage a broader perspective beyond using traditional synergy scores as the primary objective for drug combination prediction. Converting a dose-response grid into a single score inherently results in a loss of information. Such conversion oversimplifies the interaction profile of combined drugs and hides the key pharmacodynamic patterns across different dosage pairs that can be discovered across the entire dose-response surface, providing a richer representation of drug dynamics. By predicting the full dose-response surface, DeepSynBa can reveal how the effect of one drug changes in response to increasing doses of the other. More importantly, the novel intermediate layer that learns the parameters of the surface allows a clearer understanding of whether the observed interaction is driven by an enhanced potency or an increased maximal efficacy.

Future work would include making the parameters in the intermediate layer probabilistic to allow uncertainty estimation for the various aspects of the synergy. One approach is to introduce a joint distribution for these parameters and treat them as latent variables instead of fixed values, by using techniques such as the Variational Information Bottleneck (VIB) [Alemi et al., 2016]. Such a framework would provide principled uncertainty estimates, enabling prioritisation of combinations with strong synergistic potency and/or efficacy accompanied with low uncertainty for follow-up experimental validation. The wide distributions for the predicted paramaters observed in Fig. S6 further motivate this direction, as a probabilistic intermediate layer would encourage better-calibrated predictions by explicitly penalising excessive dispersion during training.

Second, we could adapt DeepSynBa for patient-level personalisation. Recent work such as PDSP [Kuru et al., 2024] has demonstrated that personalised drug sensitivity prediction from patient-derived samples is feasible and clinically relevant. Building on this, we aim to integrate patient-specific molecular and ex vivo response data into the DeepSynBa framework, so that the disentangled potency-and efficacy-related parameters can be used to propose personalised combination strategies.

Overall, DeepSynBa supports decision-making processes in drug discovery pipelines by predicting actionable drug combination response profiles. The case studies presented in this paper provide an example of how the model provides enhanced interpretability and guides the use of the combination regimes in practical applications.

