## Supplementary material for "DeepSynBa: Actionable Drug Combination Prediction with Complete Dose-Response Profiles": 6 Supplemental Tables and 6 Supplemental Figures

### Supplementary Information for DeepSynBa: Actionable Drug Combination Prediction with Complete Dose-Response Profiles

#### 1 Tables

| | **RMSE** | **Pearson** | **Spearman** |
|---|---|---|---|
| **DeepSynBa ∼ comboFM** | $4.78 \times 10^{-24}$ | $6.52 \times 10^{-33}$ | $8.18 \times 10^{-23}$ |
| **DeepSynBa ∼ comboLTR** | $3.51 \times 10^{-34}$ | $1.26 \times 10^{-30}$ | $5.60 \times 10^{-31}$ |
| **DeepSynBa ∼ comboKR 2.0** | $7.31 \times 10^{-48}$ | $2.37 \times 10^{-66}$ | $3.71 \times 10^{-58}$ |
| **DeepSynBa ∼ DD-PRISM** | $1.47 \times 10^{-30}$ | $6.66 \times 10^{-32}$ | $3.27 \times 10^{-22}$ |
| **comboFM ∼ comboLTR** | $4.35 \times 10^{-10}$ | $6.19 \times 10^{-18}$ | $1.08 \times 10^{-6}$ |
| **comboFM ∼ comboKR 2.0** | $1.04 \times 10^{-37}$ | $3.79 \times 10^{-69}$ | $6.37 \times 10^{-29}$ |
| **comboFM ∼ DD-PRISM** | $1.59 \times 10^{-1}$ | $7.92 \times 10^{-2}$ | $7.44 \times 10^{-1}$ |
| **comboLTR ∼ comboKR 2.0** | $1.04 \times 10^{-24}$ | $5.82 \times 10^{-21}$ | $3.29 \times 10^{-13}$ |
| **comboLTR ∼ DD-PRISM** | $5.00 \times 10^{-8}$ | $1.16 \times 10^{-15}$ | $6.81 \times 10^{-6}$ |
| **comboKR 2.0 ∼ DD-PRISM** | $8.45 \times 10^{-36}$ | $1.74 \times 10^{-64}$ | $8.83 \times 10^{-27}$ |

**Table S1:** The pairwise t-test $p$-value results for cell line based results in NCI-ALMANAC.

| | RMSE | Pearson | Spearman |
|---|---|---|---|
| **DeepSynBa ∼ comboFM** | $6.11 \times 10^{-38}$ | $4.17 \times 10^{-33}$ | $4.38 \times 10^{-14}$ |
| **DeepSynBa ∼ comboLTR** | $1.39 \times 10^{-55}$ | $4.63 \times 10^{-58}$ | $2.29 \times 10^{-25}$ |
| **DeepSynBa ∼ comboKR 2.0** | $4.66 \times 10^{-37}$ | $1.23 \times 10^{-44}$ | $6.58 \times 10^{-42}$ |
| **DeepSynBa ∼ DD-PRISM** | $8.85 \times 10^{-33}$ | $5.98 \times 10^{-26}$ | $3.15 \times 10^{-13}$ |
| **comboFM ∼ comboLTR** | $2.26 \times 10^{-15}$ | $1.41 \times 10^{-25}$ | $4.52 \times 10^{-4}$ |
| **comboFM ∼ comboKR 2.0** | $1.30 \times 10^{-23}$ | $3.48 \times 10^{-30}$ | $2.06 \times 10^{-13}$ |
| **comboFM ∼ DD-PRISM** | $2.73 \times 10^{-1}$ | $5.41 \times 10^{-1}$ | $9.68 \times 10^{-1}$ |
| **comboLTR ∼ comboKR 2.0** | $1.05 \times 10^{-13}$ | $2.73 \times 10^{-10}$ | $5.11 \times 10^{-5}$ |
| **comboLTR ∼ DD-PRISM** | $2.09 \times 10^{-10}$ | $7.52 \times 10^{-20}$ | $7.91 \times 10^{-4}$ |
| **comboKR 2.0 ∼ DD-PRISM** | $3.8 \times 10^{-22}$ | $1.67 \times 10^{-28}$ | $1.52 \times 10^{-12}$ |

**Table S2:** The pairwise t-test $p$-value results for drug based results in NCI-ALMANAC.

| | **Hyperparameter** | **Values** |
|---|---|---|
| Drug-Cell Encoder | hidden layers | $256, 512, 1024, 2048, 4096$ |
| | activations | ReLU |
| | input dropout | none, 0.2, $0.3, 0.5$ |
| | dropout | $0.2, 0.5$ |
| | normalizations | batch normalization |
| Prediction heads | hidden layers | $64, 128, 256$ |
| | activations | ReLU |
| | final layer activation | ReLU, sigmoid, linear |
| | normalizations | layer normalization |
| Training parameter | number of epochs | 500 |
| | loss | MSE, L1 |
| | scheduler | LinearLR |
| | optimizer | AdamW (lr=10^{-4}, wc=10^{-5}) |
| | batch size | 128 |

**Table S3:** The search space for tuning hyperparameters and training parameters for DeepSynBa architecture.

| | **MolFormer** | **ECFP4** |
|---|---|---|
| **RMSE** | **7.72** | **7.37** |
| **Pearson** | **0.95** | **0.96** |
| **Spearman** | **0.82** | **0.83** |

**Table S4:** Comparison of model performance across different drug representations. MolFormer uses transformer-based MolFormer model to extract drug features while ECFP4 is the drug fingerprints.

| | **GEX only** | **GEX + MUT + CNV + METHY** |
|---|---|---|
| **RMSE** | 7.72 | 8.66 |
| **Pearson** | 0.95 | 0.94 |
| **Spearman** | 0.82 | 0.78 |

**Table S5:** Comparison of model performance across different cell line representations. The GEX-only model relies solely on gene expression profiles, while the GEX + MUT + CNV + METHY model integrates additional genomic features, including mutations, copy number variations, and DNA methylation.

| | DeepSynBa | Matrix Prediction Head |
|---|---|---|
| RMSE | **7.72** | **22.57** |
| Pearson | **0.95** | **0.69** |
| Spearman | **0.82** | **0.54** |

**Table S6:** Comparison of alternative prediction head that directly outputs dose–response matrices. Instead of predicting SynBa parameters and reconstructing responses via the likelihood formulation, the model predicts a 4 × 4 response matrix $R_i$ for each drug in the combination. The final dose–response matrix is obtained by integrating these predicted matrices with the corresponding drug dosages.

#### 2 Figures

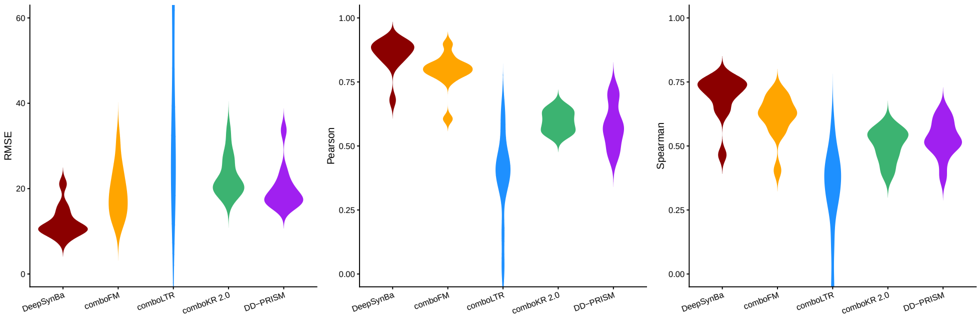


**(a)** Cell line based results on new cell scenario.

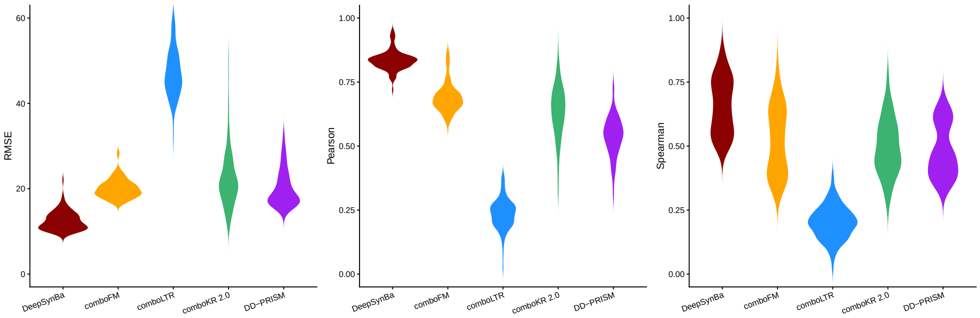


**(b)** Drug based results on new cell scenario.

**Figure S1:** The distributions of RMSE, Pearson and Spearman correlations of dose-response surface predictions on different (a) cell line and (b) drugs with new cell scenario in NCI-ALMANAC.

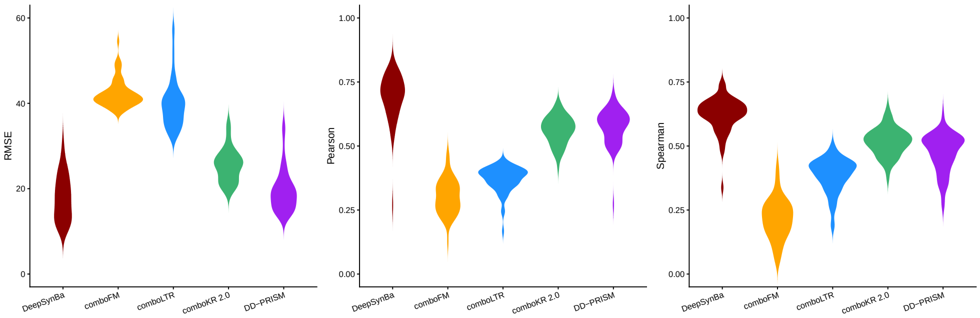


**(a)** Cell line based results on new drug scenario.

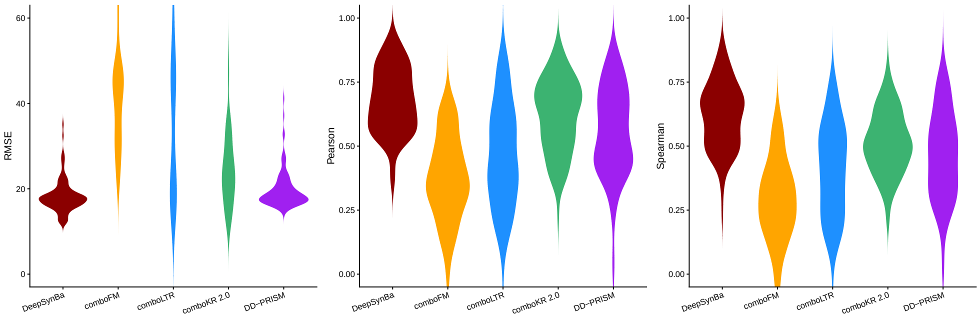


**(b)** Drug based results on new drug scenario.

**Figure S2:** The distributions of RMSE, Pearson and Spearman correlations of dose-response surface predictions on different (a) cell line and (b) drugs with new drug scenario in NCI-ALMANAC.

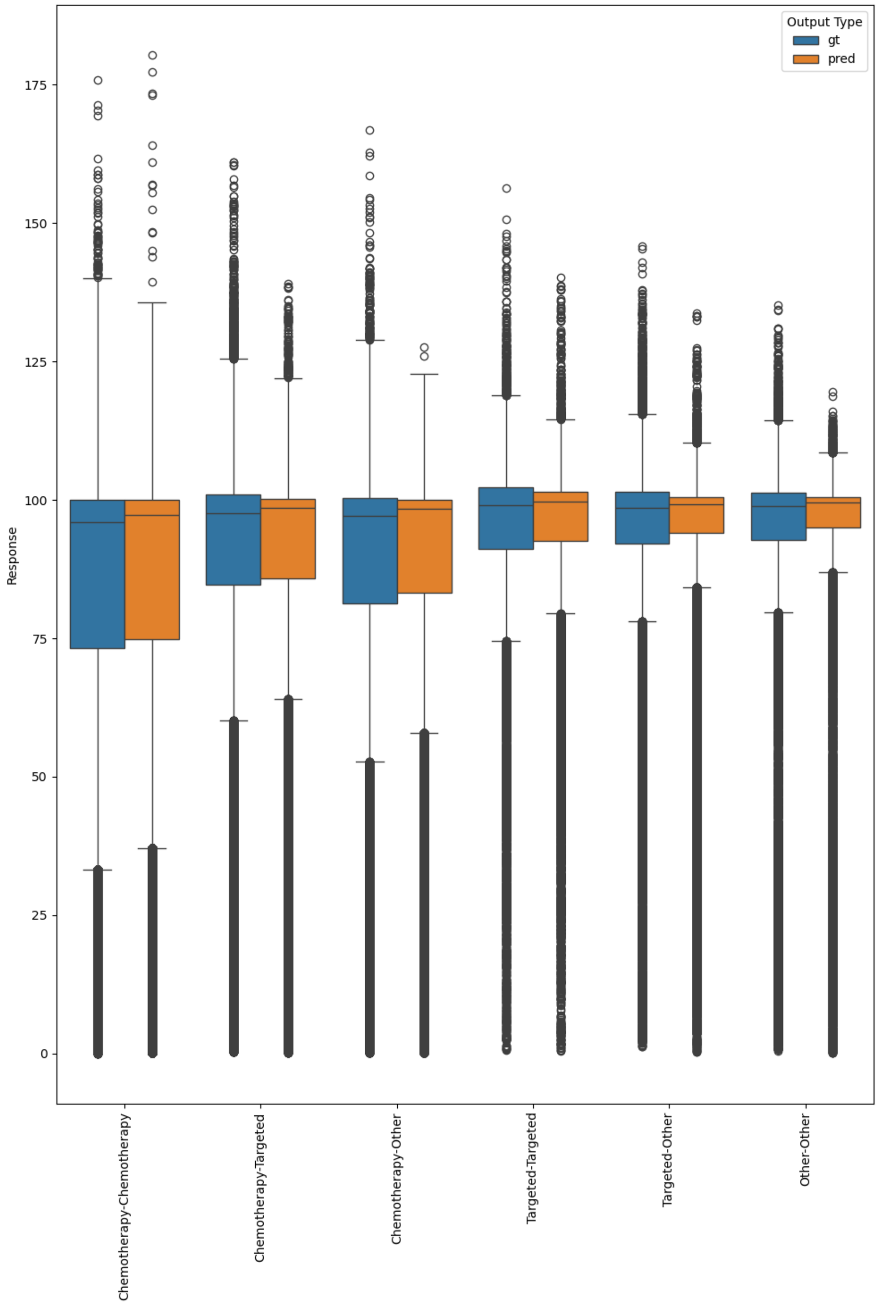


**Figure S3:** The ground-truth and the predicted response distributions for drug types in the combination for NCI-ALMANAC.

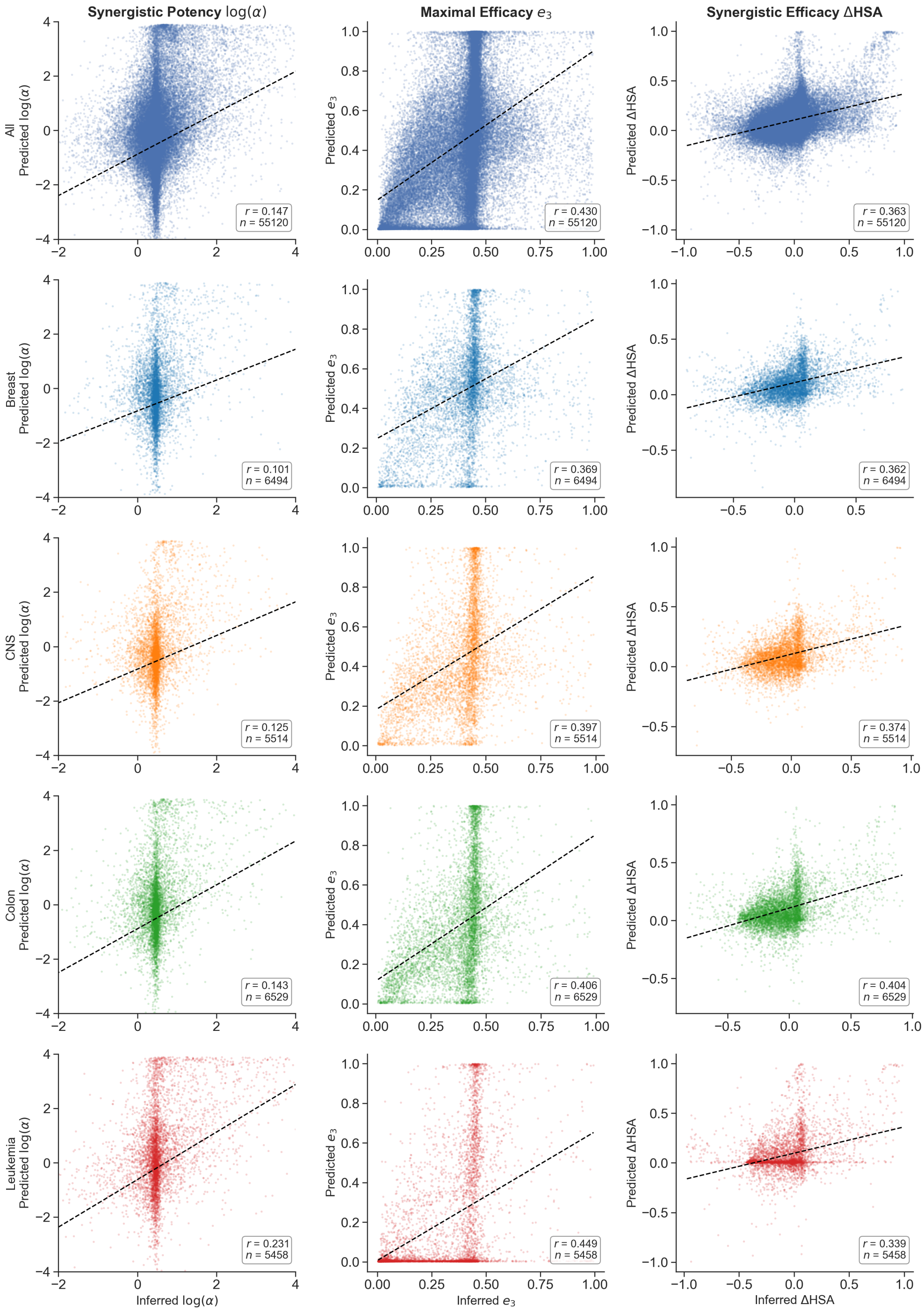


**Figure S4:** Tissue-stratified comparison of estimated and predicted synergy parameters in Scenario (i) (unseen drug combinations) of NCI-ALMANAC. The first row includes all cell types. Each subsequent row corresponds to an individual cancer tissue type. All rows show the same three metrics, i.e. (left) the synergistic potency ($\log \alpha$), (middle) the maximal efficacy ($e_3$), and (right) the synergistic efficacy ($\Delta\text{HSA} = \min\{e_1, e_2\} - e_3$). Pearson correlation coefficient $r$ and sample size $n$ are reported per panel. Dashed lines denote ordinary least-squares fits.

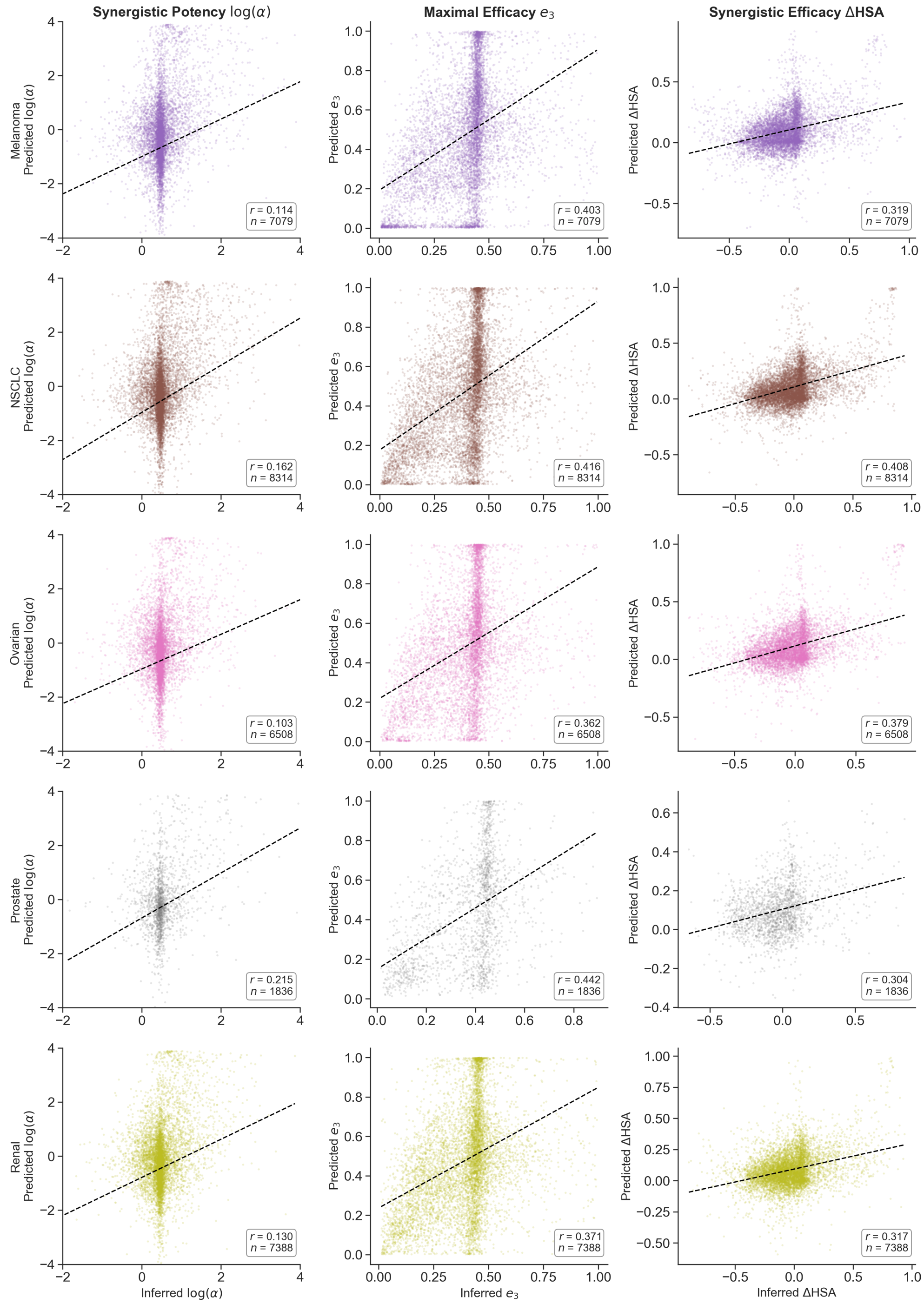


**Figure S4:** (Continued)

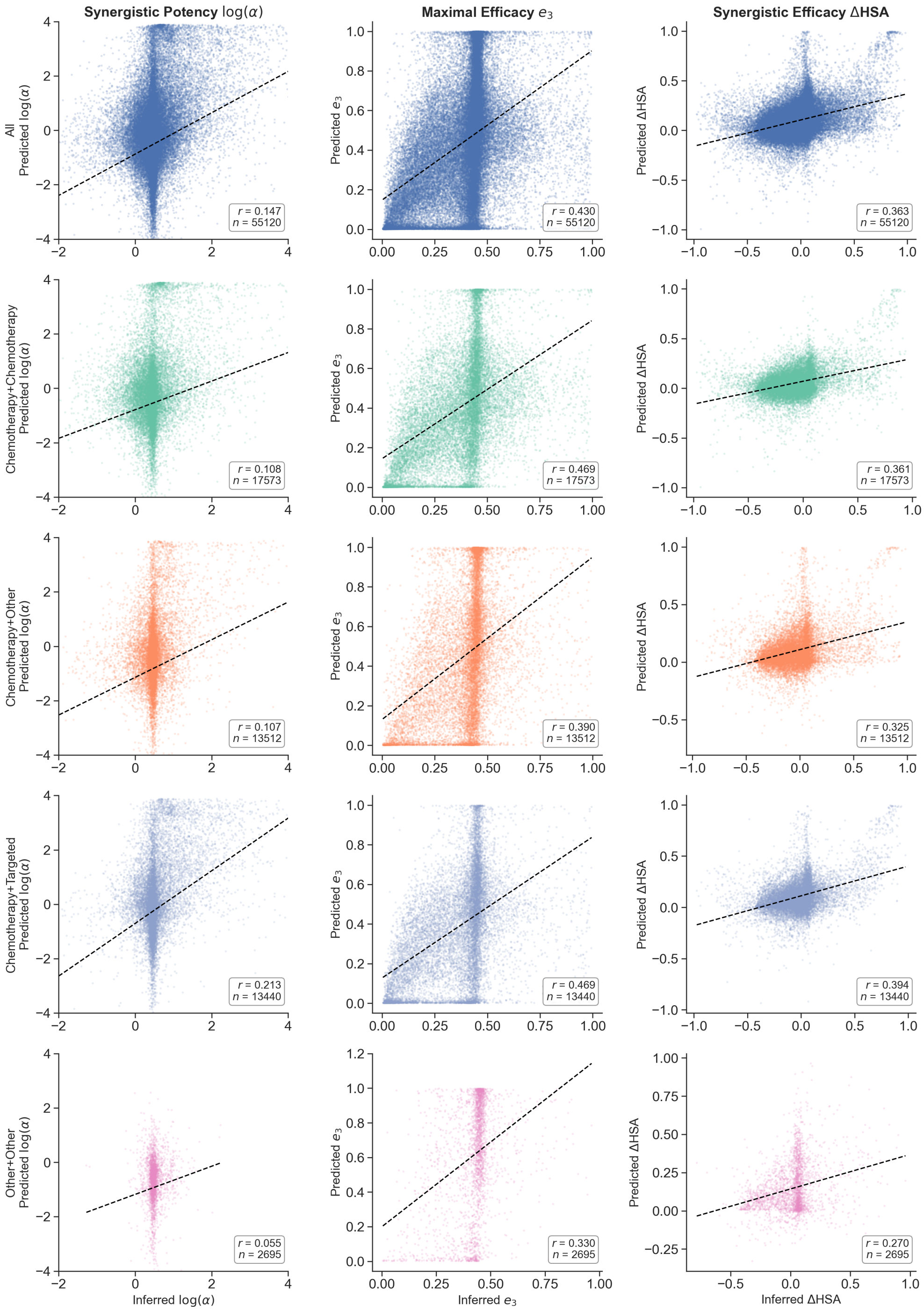


**Figure S5:** Drug-type-stratified comparison of estimated and predicted synergy parameters in Scenario (i) (unseen drug combinations) of NCI-ALMANAC. The first row includes all cell types. Each subsequent row corresponds to a pairwise combination of drugs from distinct therapeutic classes. All rows show the same three metrics as Fig. S4. Pearson correlation coefficient $r$ and sample size $n$ are reported per panel. Dashed lines denote ordinary least-squares fits.

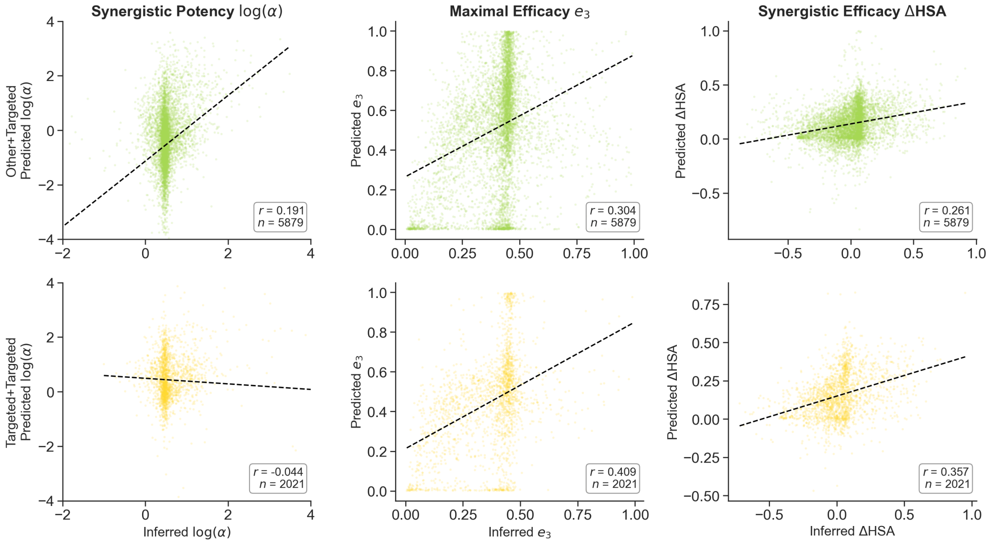


**Figure S5:** (Continued)

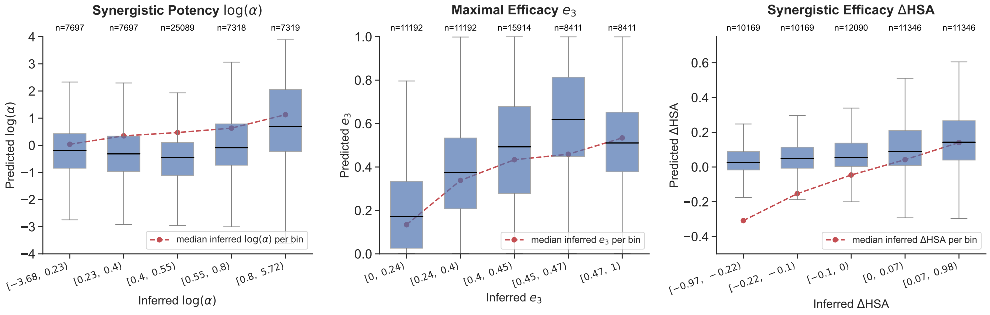


**Figure S6:** Distribution of predicted parameter values stratified by inferred value, for (left) the synergistic potency ($\log \alpha$), (middle) the maximal efficacy ($e_3$), and (right) the synergistic efficacy ($\Delta\text{HSA} = \min\{e_1, e_2\} - e_3$). For each parameter, combinations are partitioned into five bins along the inferred-value axis. The bin construction is identical across panels, consisting of a narrow central bin spanning the modal region of the inferred distribution (bin 3), and two outer bins on each side whose split points are the medians of the below-central and above-central populations, ensuring that bins 1 and 2 contain equal counts, as well as bins 4 and 5. Within each bin, the distribution of predicted values is summarised as a boxplot. The red dashed line connects, at each bin, the median of the inferred values falling in that bin. Under a perfectly calibrated model, the black median bar of each box would coincide with this marker. Sample sizes $n$ are annotated above each box.
